# Arbovirus emergence in temperate climates: the case of Córdoba, Argentina, 2009-2018

**DOI:** 10.1101/602003

**Authors:** Michael A. Robert, Daniela Tatiana Tinunin, Elisabet Marina Benitez, Francisco Ludueña-Almeida, Moory Romero, Anna M. Stewart-Ibarra, Elizabet Lilia Estallo

## Abstract

The distribution of arbovirus disease transmission is expanding from the tropics and subtropics into temperate regions worldwide. The aim of this study was to characterize the emergence of arboviruses in the temperate city of Córdoba, Argentina (2009-2018), including dengue virus (DENV) serotypes and origins of imported cases. The first cases of dengue fever were reported in 2009, followed by outbreaks in 2013, 2015, and 2016, each outbreak having greater incidence than the previous. DENV1 was the predominant serotype. Cases were imported from Venezuela, Brazil, Bolivia, Mexico, Costa Rica, and northern Argentina. The first imported cases of chikungunya were reported in 2014 and the first imported and autochthonous of Zika fever in 2016. Regional efforts are needed to strengthen surveillance, due to the key role of human movement in arbovirus introductions.

## Background

Dengue fever re-emerged in Latin America and the Caribbean in the 1980s, following the decline of widespread *Aedes aegypti* mosquito control programs aimed at eliminating Yellow Fever^1^. Dengue fever is caused by dengue virus (DENV serotypes 1-4), which causes a spectrum of acute febrile illness^2^. In Argentina, dengue was reported for the first time in over 80 years in the northwestern Province of Salta in 1997, and has since been largely constrained to northern provinces of the country with subtropical climates^3,4^.

Within the last decade, dengue emerged for the first time in areas with temperate climates, including Córdoba, the second largest city in Argentina (population 1.3 million), located in the southern cone of South America (Figure 1). The first dengue outbreak in Córdoba occurred in 2009^5^, fourteen years after *Ae. aegypti* was first detected in the city^6^. Since that time, Córdoba has reported imported and autochthonous cases of *Aedes*-transmitted arboviruses most years; however, no prior studies have characterized the local epidemiological situation. Córdoba (31.4°S, 64.2°W) is among the southernmost cities in the Western Hemisphere to report autochthonous dengue transmission, making it an important site to study the dynamics of emergence of arboviruses in southern temperate latitudes.

**Figure 1.**
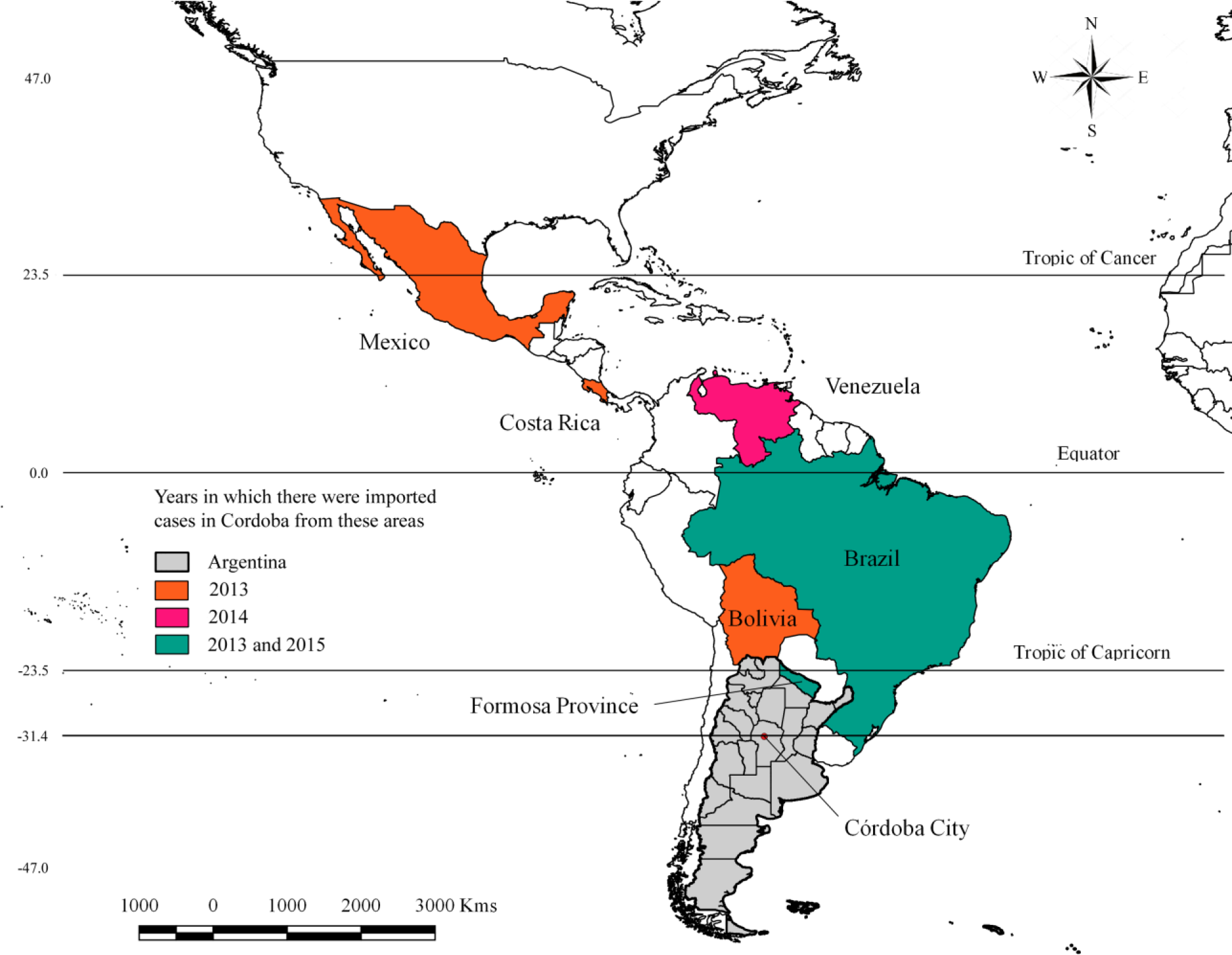
Location of Córdoba city within the province of Córdoba in Argentina. The orange, pink, and green highlighted countries are countries from which at least one dengue case was known to have been imported. Origins of imports were unknown for the majority of dengue cases reported. Lines of latitude are shown to emphasize the location of Córdoba city in relationship to the tropics.

Córdoba has a continental temperate climate, with warm summers (October – May, mean max temp = 25.1°C) and dry, cool winters (June – September, mean min temp =12.5°C). Summer temperatures fall within the lower range of optimal temperatures for arbovirus transmission by *Aedes aegypti*, which peak at 28.5°C (range: 13.5°C-34.2°C)^7^. The Ministry of Health (MoH) has reported no vector activity during winter months; therefore, local outbreaks depend on the importation of arboviruses, often by travelers from disease endemic areas.

The aim of this study was to present the epidemiological characteristics of the emergence of dengue fever and other arboviral diseases (i.e., chikungunya and Zika fever) in Córdoba over the last decade. We obtained epidemiological data by manually extracting records from weekly reports published in Spanish by the Argentinian MoH^8^. We translated, compiled, and reviewed the data to determine weekly cases, source of imported cases, and DENV serotypes in circulation. In Córdoba, epidemiological and vector surveillance are the responsibilities of the Epidemiology Area of the Zoonosis Program of the MoH of the province (M. Ainete, *pers. comm.*). Suspected dengue infections that are diagnosed at either private or public clinical sites are reported to the National Health Surveillance System operated by the MoH. A subset of clinically diagnosed cases are confirmed by laboratory diagnostics (polymerase chain reaction and immunoglobulin M antibodies to DENV) at the central MoH laboratory in Córdoba. A subset of samples are sent to the Maiztegui Institute in Buenos Aires for confirmation using the plaque reduction neutralization test (PRNT).

We consolidated all reported cases of dengue, Zika, and chikungunya for Córdoba city between January 2009 and June 2018 (114 weekly reports). We cleaned the data when inconsistencies were noted (e.g., eliminating cases that appeared to be counted multiple times). We aimed to collect all available data on arbovirus cases including probable (clinically diagnosed) and laboratory confirmed cases, autochthonous (locally transmitted) and imported cases (illness in someone with a travel history), DENV serotypes, and origins of imported cases. However, this information was not available in all reports. The cases presented here are a sum of probable and confirmed cases unless otherwise noted. Herein, we present a synthesis of arbovirus transmission in Córdoba city since 2009, and focus on the characteristics of the four major outbreaks of dengue that occurred between 2009 and 2018.

## Arbovirus outbreaks

In Figure 2, we present weekly incidence of cases from January 2009 to June 2018. A total of 1,429 dengue cases were reported during this period (1,170 autochthonous, 259 imported). DENV1 was the predominant serotype in circulation over the last decade, and DENV4 played a secondary role, although all four DENV serotypes were detected (Table 1). Imported dengue cases originated from tropical countries where dengue fever is endemic, as well as the endemic subtropical northern region of Argentina.

**Table 1.** Dengue virus serotypes and origin of imported dengue cases. N is the number of cases in Córdoba that were tested for serotype each year.

| Year | DENV Serotypes Detected | Origins of Imported Cases |
| --- | --- | --- |
| 2009 | DENV-1 (Majority; N unknown) | No Information |
| 2010 | No Information |  |
| 2011 | No Information |  |
| 2012 | DENV-1 (50%), DENV-4 (50%); (N=2) | No Information |
| 2013 | DENV-1 (73.8%), DENV-2 (1.6%), DENV-3 (3.3%),<br>DENV-4 (21.3%); (N=61) | Brazil, Bolivia, Mexico,<br>Costa Rica, Formosa<br>Province (Argentina) |
| 2014 | DENV-1 (100%); (N=2) | Venezuela |
| 2015 | DENV-1, DENV-4 (Frequency Unknown) | Brazil, Formosa Province<br>(Argentina) |
| 2016 | DENV-1 (98.6%), DENV-2 (0.4%), DENV-3 (0.4%),<br>DENV-4 (1.0%); (N=288). | No Information |
| 2017 | No Information |  |
| 2018 | DENV-1 (Majority; N Unknown) | Colombia |

**Figure 2.**
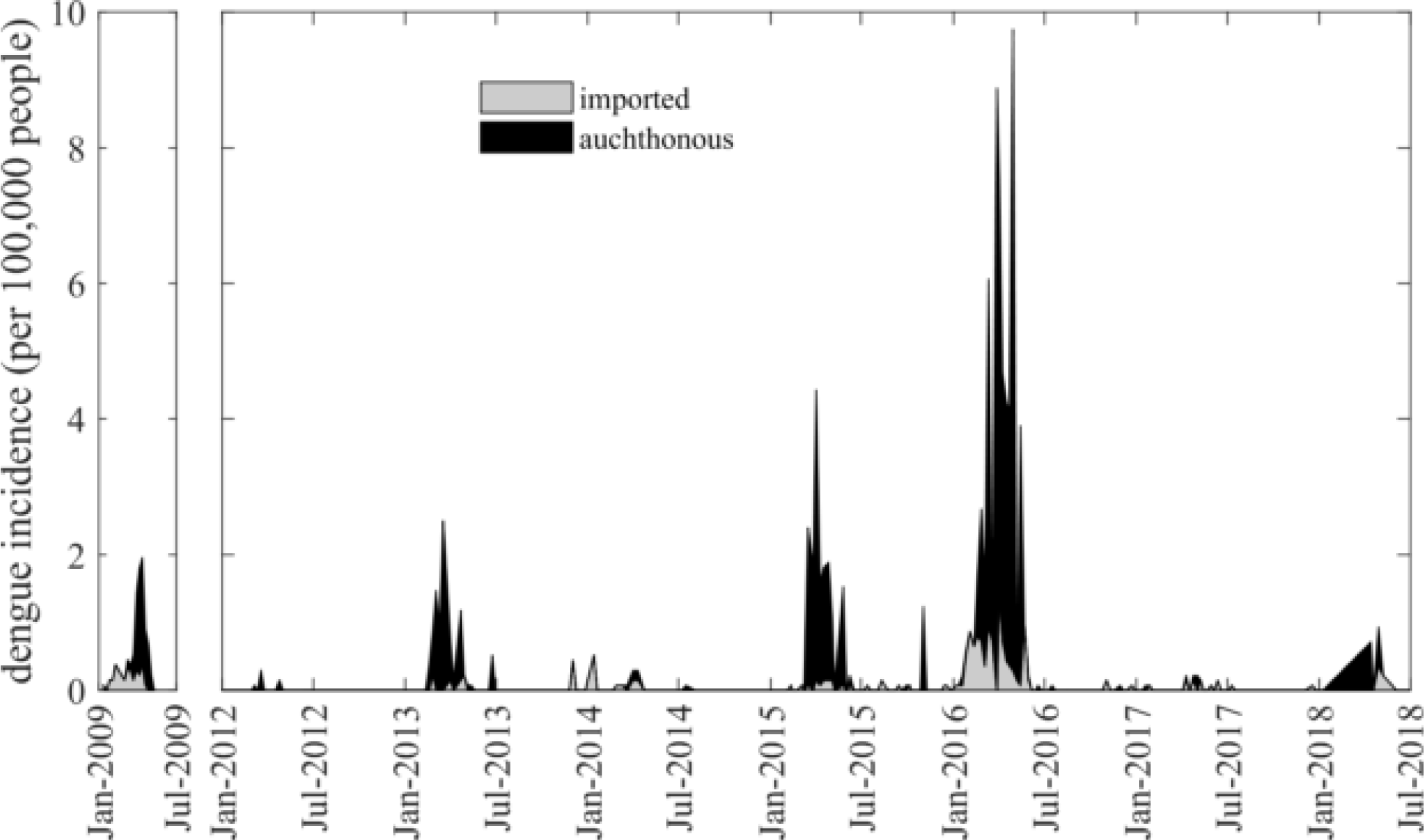
Incidence of imported (gray) and autochthonous (black) dengue cases relative to total incidence each epidemic week between January 2009-July 2018. Incidence is calculated as the number of cases per 100,000 inhabitants of Córdoba, where yearly population was estimated using census data from 2001^13^, 2008^14^, and 2010^8^, and linear interpolation. The population size as estimated by the function P(t) = 8045.47t + 1257194.75 and t=0 is 2001^15^. Note: there was no reported dengue activity in 2010-2011, so this period has been excluded from the figure.

The first imported dengue case in Córdoba was reported Epidemiological Week (EW) 2 in 2009. In EW10, 2 cases of dengue in patients without travel to affected areas were confirmed, marking the first known autochthonous dengue case in Córdoba. In total, 88 autochthonous cases were reported between EW10-EW18. During EW2-EW18, 42 imported dengue cases were reported, leading to 130 total cases. All tested cases were confirmed to be DENV1^9^. The total dengue incidence in 2009 was 9.78 cases per 100,000 people.

In 2013, Córdoba experienced its second major outbreak, with 115 autochthonous and 10 imported cases reported (total incidence 10.13). Autochthonous and imported cases were reported from EW7-18. The known origins of imported cases include Brazil, Bolivia, Mexico, Costa Rica, and Formosa Province in northeastern Argentina.

In 2015, Córdoba experienced its third significant outbreak of dengue beginning with imported cases in EW5. From EW9-22, 221 autochthonous cases were reported. In total, 236 autochthonous and 14 imported dengue cases were reported (total incidence 19.01). DENV1 and DENV4 were detected, and cases were imported from Brazil and Formosa Province, Argentina.

Córdoba’s largest dengue outbreak to date began in EW52 of 2015 with an imported case of unknown origin. The first 2 autochthonous cases were reported in EW2, and 687 autochthonous and 134 imported cases were reported from EW2-24. In total, 822 cases of dengue were reported (688 autochthonous; 134 imported; total incidence 60.25). Of these, 288 (35%) cases were tested for DENV serotypes (Table 1). DENV2, DENV3, and DENV4 serotypes were detected in imported cases. Of the 284 DENV1 cases, 221 (78%) were autochthonous, and 63 (22%) were imported.

Chikungunya and Zika fever first emerged in Córdoba in 2014 and 2016, respectively (Table 1, Supplementary Material). A total of 22 imported chikungunya cases were reported from 2014 to 2017. In 2016, 4 imported and 1 autochthonous cases of Zika were reported. No cases of congenital Zika syndrome were detected.

## Discussion

In the last decade, *Aedes*-transmitted arbovirus-related illness has emerged for the first time in the city of Córdoba. The city has quickly become a site of significant epidemic dengue transmission, as indicated by the occurrence of 4 outbreaks in less than 10 years, and local transmission most years since 2009. During outbreaks, cases peaked during the later part of the summer, when vector densities were elevated and maximum daily temperatures were slightly lower than the optimum temperature for arbovirus transmission by *Aedes aegypti* of 28.5°C^7^. The increase in arbovirus activity in Córdoba mirrors that of arbovirus activity across temperate regions of the world, including the southern United States and southern Europe^9^. Autochthonous cases of dengue re-emerged after many decades of no transmission in Florida, USA, in 2009 ^10^, and in France, Croatia, and Portugal in 2010-2013^11^. Increasing arbovirus transmission in temperate latitudes is likely associated with social and ecological factors including greater human movement, expansion and local adaptation of *Aedes* mosquitoes, and changes in climate resulting in increased surface temperatures and altered rainfall patterns^12^.

As Córdoba is a temperate region, local transmission depends on the importation of cases from dengue endemic areas. With the current political crisis in Venezuela, resulting in mass migration of people into Argentina, the risk of importation of dengue fever and other mosquito borne diseases (e.g., malaria) has increased^13^. Regional efforts are needed to strengthen surveillance of arbovirus transmission, due to the key role of human movement in dengue introductions, as shown here.

In response to the emergence of dengue fever, the MoH of Argentina has implemented a comprehensive *Ae. aegypti* surveillance program across Córdoba using larval surveys and ovitraps during the season of vector activity in cooperation with the Córdoba Entomological Research Centre from the National University of Córdoba and the Institute of Biological Research (IIBYT). Vector control by the MoH is mostly focal, around homes with dengue cases, and includes control of adult mosquito populations (indoor and outdoor fumigation) and larval mosquitoes (*Bacillus thuringiensis israelensis* larvicide application, elimination of larval habitat) ^14^. However, these efforts were unable to prevent outbreaks (as in 2016), and risk perception by the public remains low. Further investigation of the role of social and environmental drivers of arbovirus emergence in Córdoba is needed to develop effective vector control and disease management programs to reduce the burden of dengue illness.

## Acknowledgements

This work was supported by grants awarded to A.M. Stewart-Ibarra and M.A. Robert by the United States Embassy in Argentina administered through the Fulbright Commission. ELE is a member of the Consejo de Investigaciones Cientificas y Tecnologicas (CONICET) from Argentina.

## Supplemental Material

**Table 1.** Imported and autochthonous chikungunya and Zika virus cases in Córdoba.

| Year | Chikungunya |  | Zika Virus |  |
| --- | --- | --- | --- | --- |
|  | Imported | Autochthonous | Imported | Autochthonous |
| 2014 | 10 | 0 |  |  |
| 2015 | 4 | 0 |  |  |
| 2016 | 6 | 0 | 4 | 1 |
| 2017 | 2 | 0 | 0 | 0 |
| 2018 | 0 | 0 | 0 | 0 |

## References

1. Gubler DJ, Clark GG. Dengue/dengue hemorrhagic fever: the emergence of a global health problem. Emerg Infect Dis. 1995;1(2):55–57.

2. WHO. Dengue: Guidelines for Diagnosis, Treatment, Prevention and Control. World Health Organization; 2009.

3. Vezzani D, Carbajo AE. Aedes aegypti, Aedes albopictus, and dengue in Argentina: current knowledge and future directions. Memórias do Instituto Oswaldo Cruz. 2008;103(1):66–74.

4. Aviles G, Rangeón G, Vorndam V, Briones A, Baroni P, Enria D, Sabattini MS. Dengue reemergence in Argentina. Emerging Infectious Diseases. 1999;5(4):575.

5. Estallo EL, Carbajo AE, Grech MG, Frías-Céspedes M, López L, Lanfri MA, Ludueña-Almeida FF, Almirón WR. Spatio-temporal dynamics of dengue 2009 outbreak in Córdoba City, Argentina. Acta Tropica. doi:10.1016/j.actatropica.2014.04.024.

6. Almirón WR, Almeida FL. Aedes aegypti (Diptera: Culicidae) en Córdoba, Argentina. Revista de la Sociedad Entomológica Argentina. 1998;57(1-4).

7. Mordecai EA, Cohen JM, Evans MV, Gudapati P, Johnson LR, Lippi CA, Miazgowicz K, Murdock CC, Rohr JR, Ryan SJ, others. Detecting the impact of temperature on transmission of Zika, dengue, and chikungunya using mechanistic models. PLoS Neglected Tropical Diseases. 2017;11(4):e0005568.

8. Ministerio de Salud de Nación (MSN). Boletines Epidemiologicos. https://www.argentina.gob.ar/salud/epidemiologia/boletinesepidemiologicos. Accessed April 28, 2018.

9. Messina JP, Brady OJ, Scott TW, Zou C, Pigott DM, Duda KA, Bhatt S, Katzelnick L, Howes RE, Battle KE, Simmons CP, Hay SI. Global spread of dengue virus types: mapping the 70 year history. Trends in Microbiology. 2014;22(3):138–146. doi:10.1016/j.tim.2013.12.011.

10. Radke EG, Gregory CJ, Kintziger KW, Sauber-Schatz EK, Hunsperger EA, Gallagher GR, Barber JM, Biggerstaff BJ, Stanek DR, Tomashek KM. Dengue outbreak in key west, Florida, USA, 2009. Emerging infectious diseases. 2012;18(1):135.

11. Dengue and dengue vectors in the WHO European region: past, present, and scenarios for the future - The Lancet Infectious Diseases. https://www.thelancet.com/journals/laninf/article/PIIS1473-3099(14)70834-5/fulltext. Accessed February 21, 2019.

12. Muñoz ÁG, Thomson MC, Goddard L, Aldighieri S. Analyzing climate variations at multiple timescales can guide Zika virus response measures. Gigascience. 2016;5(1):41.

13. Jaramillo R, Sippy R, et al. Ahead of Print - Effects of Political Instability in Venezuela on Malaria Resurgence at Ecuador–Peru Border, 2018 - Volume 25, Number 4—April 2019 - Emerging Infectious Diseases journal - CDC. doi:10.3201/eid2504.181355.

14. PLAN NACIONAL PARA LA PREVENCIÓN Y CONTROL DEL DENGUE Y LA FIEBRE AMARILLA. July 2009. http://www.msal.gob.ar/images/stories/cofesa/2009/acta-02-09/anexo-5-resumen-plan-dengue-02-09.pdf. Accessed April 25, 2018.

15. Dirección general de estadistica y censos, Gobierno de la Provincia de Córdoba. Censo Provincial de Población 2008: Municipio de Córdoba. Censo 08 de Poblacion de la Provincia Córdoba.

